# Monitoring cancer evolution under iatrogenic selection pressure by cellular liquid biopsy

**DOI:** 10.64898/2026.11.13.766382

**Authors:** Clara Chaiban, Jonas Roth, Adithi Ravikumar Varadarajan, Jonas Grote, Vadim Dechand, Thomas Ragg, Robin Gähr, Kirsten Utpatel, Wolfgang Dietmaier, Stephan Seitz, Jens Warfsmann, Christoph A. Klein, Zbigniew T. Czyż

## Abstract

Cell-free liquid biopsy (LB) provides information about real-time tumor burden in cancer patients. In contrast, cellular LB can resolve the co-occurrence of actionable mutations within individual CTCs and their clonal architecture, revealing emerging treatment opportunities, information increasingly needed as the number of available drugs grows. Here, we describe a case with repeated blood sampling and CTC analysis using a single-cell-adapted IMPACT assay (scIMPACT). We observed therapeutically relevant alterations and clonal shifts arising in response to selected therapies. Applied systematically, such analyses have the power to uncover cellular adaptations to therapy-induced selection pressures and enable flexible treatment adjustments to evolving disease.

## Main Text

Precision oncology is rapidly changing. Alone for advanced HR+/HER-breast cancer, the number of therapies increased almost exponentially over the last decade ^1^. This raises questions about how to monitor drug responses and how to change treatments best upon disease progression. While the FDA-approved CellSearch assay used to monitor circulating tumor cells (CTC) has unquestioned prognostic power ^2,3^, its adaptation to a predictive and a research tool has not been explored sufficiently. However, we previously showed that detailed molecular assessment of CTC enriched by CellSearch is feasible with diagnostic precision ^4^. We show that combining CellSearch with a single-cell adaptation of the FDA-authorized MSK-IMPACT assay ^5^ enables molecular profiling of CTCs and detection of therapeutically relevant alterations during systemic therapy, extending an approach that has already been successfully applied to FFPE-derived DNA from breast cancer specimens to characterize genomic landscapes and identify alterations associated with resistance to endocrine therapy ^6,7^.

To identify therapeutic options after rapid post-relapse progression, we explored cellular LB because (i) blood sampling is minimally invasive and can be performed repeatedly as needed; (ii) rising CTC counts under therapy can precede imaging-documented progression by 6 months ^8^, enabling early detection of relapse; and (iii) unlike circulating tumor DNA (ctDNA) analysis, cellular LB resolves the heterogeneity of disease-driving clones and emerging escape variants at cellular level. To assess clonal dynamics across treatment, we compared two baseline populations, residual PT tissue and DCCs from bone marrow (BM-DCC) aspirates obtained at surgery after neoadjuvant therapy, with samples collected after clinical manifestation of metastatic disease (hereafter, manifest metastasis [MM] samples), comprising CTCs from blood and DCCs from a malignant pleural effusion (PE-DCCs). Baseline and MM samples therefore differed in anatomical origin and in cumulative exposure to systemic therapy.

We processed 38 and analyzed 35 samples passing quality control, collected longitudinally from multiple anatomical sites of a 32-year-old premenopausal woman with luminal B breast cancer who relapsed during adjuvant abemaciclib therapy (Fig.1A,B, Table S1). PT typing revealed a high-grade, mismatch repair-proficient tumor (ER-positive, PR-negative, HER2-low, Ki-67 85%; cT2N2M0; Fig.1C,E). The patient received neoadjuvant epirubicin/cyclophosphamide followed by paclitaxel, surgery without pathological complete response, and adjuvant ovarian suppression (goserelin), letrozole and abemaciclib (Fig.1A). Despite guideline-concordant multimodal treatment, liver metastases developed within 14 months of diagnosis, during ongoing adjuvant therapy, followed by a malignant pleural effusion (PE) two months later (Fig.1A,F, Table S1). The liver metastasis lacked detectable hormone receptor expression. However, since the biopsy contained only approximately 50 tumor cells, conversion to a triple-negative phenotype should be interpreted cautiously.

**Figure 1.**
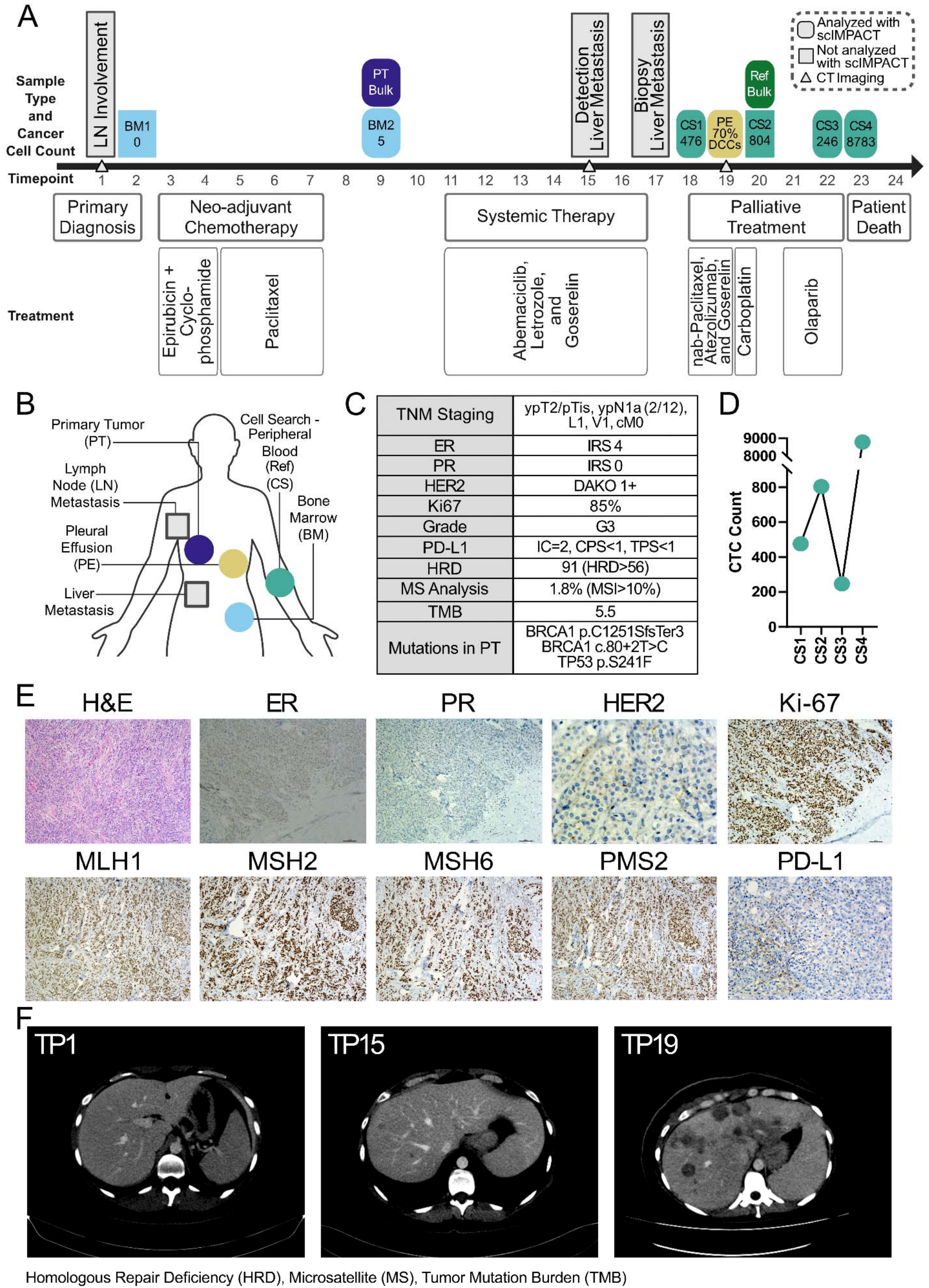
Clinical course and longitudinal single-cell sampling strategy. **(A)** Timeline of the patient’s clinical course, systemic treatments, and longitudinal sample collection, including tumor cell counts. Sample types are color-coded as shown in (B). **(B)** Overview of solid tumor and cellular LB samples detected (square) and analyzed (circles) in this study. Reference (Ref) genomic DNA was isolated from peripheral blood mononuclear cells collected at the second CellSearch sampling (CS2); **(C)** Clinicopathological and molecular characteristics of the PT; **(D)** Longitudinal CTC counts per 7.5 mL of peripheral blood, measured at TP18, TP20, TP22 and TP23 using the CellSearch® system; (E) Histopathological and immunohistochemical characterization of the PT, including H&E staining and immunostaining for ER, PR, HER2, Ki-67, MLH1, MSH2, MSH6, PMS2 and PD-L1; **(F)** Serial computed tomography (CT) images illustrating the development of liver metastases.

Longitudinal liquid biopsy comprised two BM aspirates, four peripheral blood samples analyzed with the CellSearch system (CS1–CS4) and one PE sample (Fig.1A,B, Table S1). Individual DCCs and CTCs underwent whole-genome amplification followed by scIMPACT, a single-cell adaptation of FDA-authorized MSK-IMPACT. Bioinformatic methods, quality-control criteria, and assay limitations are described in the Supplementary Methods. No BM-derived DCCs (BM-DCCs) were detected before neoadjuvant therapy, whereas five BM-DCCs were identified at surgery, indicating minimal residual disease persisting despite neoadjuvant chemotherapy. After relapse, the patient received carboplatin (between TP19 and TP20) followed by olaparib (between TP21 and TP22), which was discontinued because of adverse effects (Fig.1A). CTC counts were very high after detection of metastasis, declined during olaparib therapy and rose sharply after its discontinuation, paralleling clinical progression (Fig.1A,D).

Routine molecular profiling of the PT identified two somatic pathogenic BRCA1 alterations (BRCA1 c.80+2T>C and p.C1251Sfs^*^3), a TP53 p.S241F mutation, microsatellite stability, low tumor mutational burden and a high homologous recombination deficiency (HRD) score (Fig.1C). Although germline status was not confirmed, the molecular tumor board considered the BRCA1 findings and high HRD score sufficient to support carboplatin and subsequently olaparib. Genome-wide copy-number alteration (CNA) profiling revealed a remarkably stable backbone of chromosomal aberrations across PT and MM samples (Fig.2A,B, Fig.S1). In contrast, BM-DCCs lacked this backbone (Fig.S2) and exhibited a significantly lower percentage of aberrant genome (PAG) than all other tumor compartments (median 3.67% vs 51.16%; P < 0.0001, Mann-Whitney test; Fig.2C). The shared CNA backbone comprised recurrent gains involving *CDK6, CCNE2, MYC, CCND1* and *AURKA*, together with recurrent losses affecting *CDKN2A, PTEN, FAT1, CDK4* and *TP53* (Fig.2A,B, Fig.S1, Table S2), most of which were absent from BM-DCCs (Fig.S2). A notable exception was the *CCNE1* gain, which was absent from the PT but detected in one BM-DCC, seven CS1 CTCs, all four PE-DCCs, five CS3 CTCs and five CS4 CTCs (Fig.2B). Given the debated clinical relevance of cyclin E overexpression and *CCNE1* amplification in breast cancer, particularly for CDK4/6i resistance ^9,10^, this finding indicates that a *CCNE1*-gained clone with a potential selective advantage was present outside the PT by the end of neoadjuvant therapy and was subsequently enriched in metastatic disease.

In contrast to the stable CNA backbone, single-nucleotide variant (SNV) profiles revealed pronounced temporal and intercellular heterogeneity (Fig.2D, Table S3). BM-derived DCCs displayed the highest mutational burden, with a median of 30 medium- and high-confidence SNVs per cell (Supplementary Methods), compared with a median of 10 SNVs in MM samples (Fig. 2D). This finding indicates substantial SNV-level heterogeneity despite the comparatively low CNA burden of BM-derived DCCs relative to the other sample types. The non-linear relationship between local and systemic disease (baseline and MM samples) was further underscored by the absence of the pathogenic *BRCA1* p.C1251Sfs^*^3 mutation from all samples except the PT, despite adequate coverage of the locus (Fig.2E, Fig.S3, Table S3). The *BRCA1* c.80+2T>C variant could not be assessed in single cells (Fig.2E, Fig.S3) due to the representation bias introduced by the whole-genome amplification (WGA) used (Fig.S4, Table S4, Supplementary Methods). However, DICER1 p.F1867=, JAK1 p.V331I, ARID2 p.I1401T, MITF p.A38D, and E2F3 p.R305^*^ were detected in subsets of BM-DCCs and MM samples but not in the PT, supporting a closer relationship of BM-DCCs to MM samples than to the PT (Fig. 2E, Table S3). In contrast, TP53 p.S241F, NOTCH3 p.P2101S, and NSD1 p.G2155E were present in the PT and MM samples but were absent from BM-DCCs. Overall, the data support common descent followed by branched evolution (Fig.2E, Fig.S3, Table S3). However, neither CNA nor SNV patterns were consistent with linear transmission from PT to MM cells, from PT to BM-DCCs or from BM-DCCs to MM cells. These findings suggest that linear models of local-to-systemic cancer progression oversimplify the underlying clonal dynamics ^11^ and that the pool of potentially progression-driving variants could be larger than previously thought.

**Figure 2.**
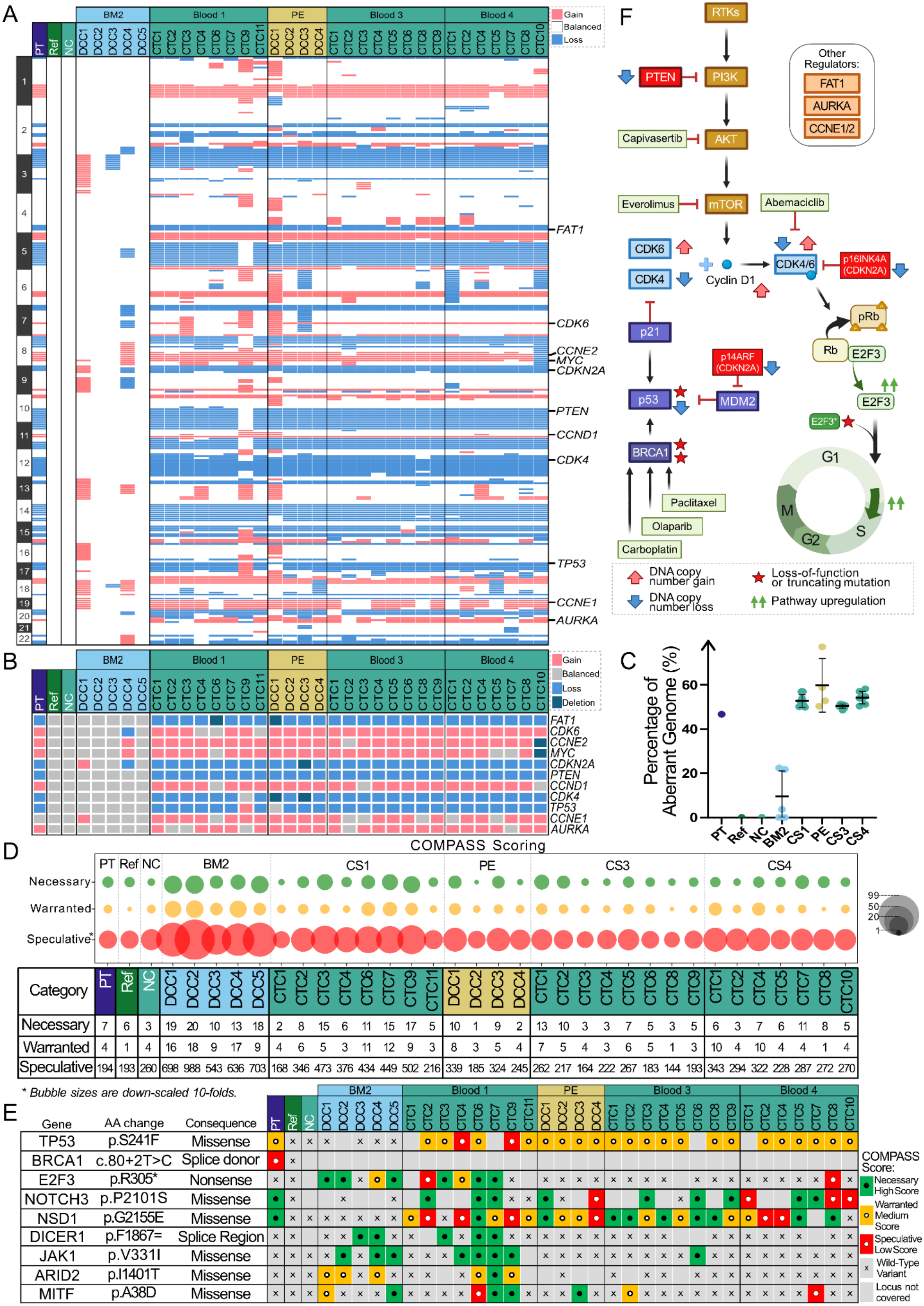
Stable copy-number architecture and dynamic SNV-based subclonal diversification define clonal evolution of the metastatic lineage. **(A)** Genome-wide CNA profiles averaged over 10-Mb genomic bins in the analyzed sample collective; **(B)** Oncoplot summarizing the frequency of CNAs involving key cell-cycle regulators and genes implicated in resistance to CDK4/6i; **(C)** Percentage of aberrant genome (PAG) in individual tumor cells, grouped by sampling timepoint; (D) Number and confidence classification of SNVs detected in individual tumor cells. Bubble size is proportional to the number of variants, with speculative-confidence variants displayed at one-tenth scale; **(E)** Distribution of selected clinically relevant SNVs and their confidence classification. Empty gray squares indicate locus dropout, gray squares marked with an × indicate wild-type calls, and colored symbols denote high-, medium- and low-confidence variants according to scIMPACT COMPASS categories - Necessary, Warranted and Speculative, respectively; **(F)** Schematic summary of recurrent genomic alterations affecting cell-cycle regulation and oncogenic signaling pathways.

In addition to the systemic selection of the *CCNE1* gain, a truncating *E2F3* p.R305^*^ mutation apparently was selected by therapy exposure. It was detected with high confidence in four of five BM-DCCs and four of eight CTCs from the first metastatic sampling (CS1) but not in subsequent samples (Fig.2E, Fig.S3, Table S3). Because this mutation leads to the truncation of the RB1-binding domain of E2F3, it may enable RB-independent G1/S transition in the presence of CDK4/6i, which currently lacks functional validation. However, its disappearance after cessation of abemaciclib is consistent with clonal fluctuation under changing treatment pressure.

Together, PT and longitudinal profiling provided information that can be linked to treatment decisions: First, the pathogenic BRCA1 alterations and high HRD score identified in the PT provided the rationale for treatment with olaparib, which was followed by a transient decline in CTC burden (Fig.1D). Second, single-cell profiling revealed expansion of *CCNE1* gain across MM samples and the temporally restricted *E2F3* p.R305^*^-mutated escape subpopulation responded to drug exposure and withdrawal (Fig.2E). However, since the single-cell analyses were performed retrospectively, they did not inform treatment in this patient.

Cellular LB captured further genomic features associated with therapy resistance. Persistent losses of *FAT1, CDKN2A, PTEN* and *TP53* together with recurrent gains involving *CDK6, CCNE2, MYC, CCND1, CCNE1* and *AURKA*, collectively indicate disruption of interconnected cell-cycle regulatory pathways (Fig.2F). Experimental and clinical studies have implicated gains of *CDK6, CCNE1, CCNE2, MYC* and *AURKA*, together with losses of *FAT1, PTEN* and *CDKN2A*, in intrinsic or acquired CDK4/6i resistance through compensatory cell-cycle programs that bypass RB-dependent G1/S control (Fig. 2F) ^12,13^. Likewise, pathogenic *TP53* and *BRCA1/2* mutations have been associated with poor responsiveness to CDK4/6 inhibition ^14–16^. Thus, scIMPACT extended routine PT profiling by revealing the coexistence of multiple resistance-associated alterations, consistent with a multifactorial basis for reduced sensitivity to CDK4/6 inhibition.

Detecting such resistance-associated genotypes during therapy could inform treatment adaptation. Preclinical studies support combined targeting of mTOR and CDK4/6 signaling ^17^, whereas the CAPItello-291 trial demonstrated that the pan-AKT inhibitor capivasertib plus fulvestrant improved progression-free survival in advanced HR+ breast cancer. Moreover, evidence suggests that *CCNE1* amplification confers sensitivity to selective CDK2 inhibition ^18,19^, while simultaneous inhibition of CDK2 and CDK4/6 has demonstrated synergistic antitumor activity in preclinical models ^20,21^. Finally, AURKA inhibition is a potential strategy for CDK4/6i-resistant tumors ^22^.

Despite the limitations of our study, including restricted genomic coverage imposed by deterministic WGA, limited functional validation of escape mechanisms and the (current) lack of therapeutic alternatives, we note several potential advantages of cellular LB combined with scIMPACT: (i) early detection of selected variants may enable rapid treatment adaptation the more targeted drugs become available; and (ii) co-occurrence of alterations associated with resistance or therapeutic response can be resolved by single-cell analysis but not by conventional ctDNA profiling. Until effective therapies exist for the full spectrum of emerging resistance mechanisms, the greatest value of this approach lies in real-time monitoring of genomic evolution under therapy, which may reveal escape mechanisms. The tools are available now. Applied at scale, they may provide fundamental insights into cancer evolution and, ultimately, enable precise treatment selection for individual patients.

## Ethics statement

The patient, treated at the Comprehensive Cancer Center Regensburg, provided written informed consent for comprehensive molecular profiling of primary tumor tissue, CTCs, DCCs, pleural effusion specimens and matched germline controls, and for review of clinical data, including pathology and treatment history. The study was approved by the ethics committee at the University of Regensburg (approval no. 18-948-101) and conducted in accordance with the Declaration of Helsinki.

**Figure S1.**
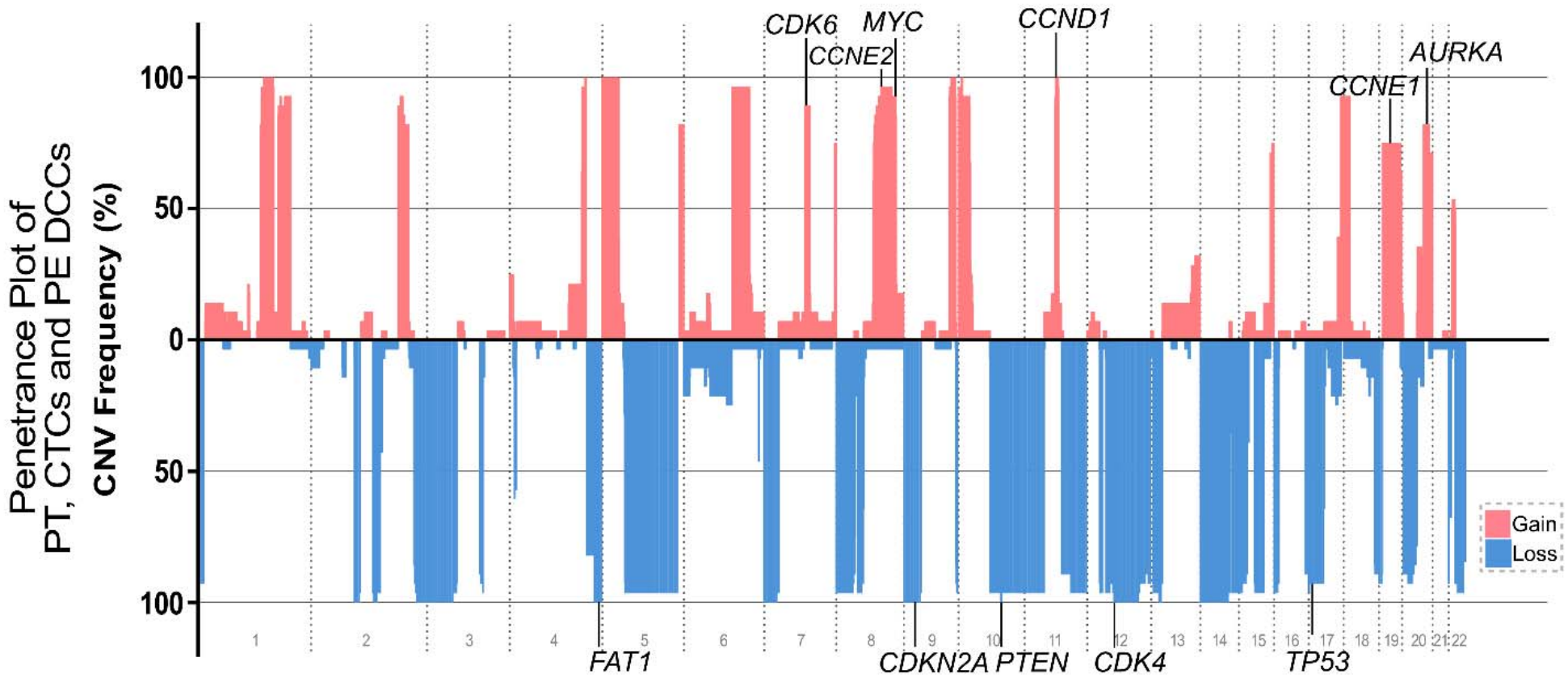
Penetrance plot of recurrent CNAs in the PT and MM samples. Frequency of recurrent CNAs across analyzed tumor sample collective, excluding BM-derived DCCs. Highlighted genes are involved in cell-cycle regulation and have been implicated in resistance to CDK4/6i.

**Figure S2.**
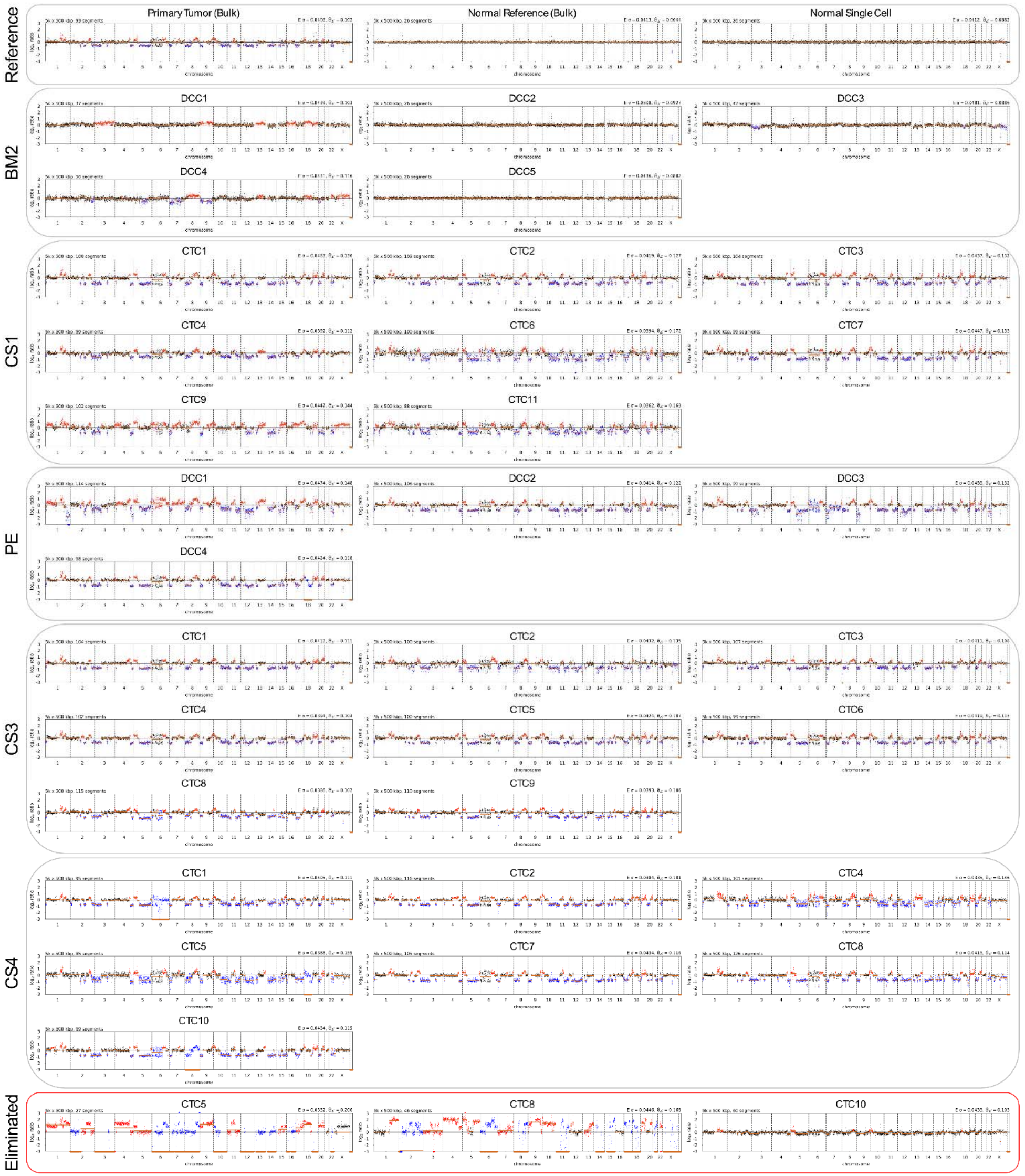
Genome-wide CNA profiles of all 38 processed samples. Horizontal CNA profiles across chromosomes 1-22 are displayed for all analyzed samples, grouped by the sampling timepoint. Log2 values on y-axis indicate copy-number gains and losses, with algorithm-defined gains, losses, and balanced regions shown in red, blue, and black, respectively. Orange lines indicate average log2 values computed for algorithm-defined genomic segments. Three samples from the first blood draw (CS1) failing quality control are marked by a red box.

**Figure S3.**
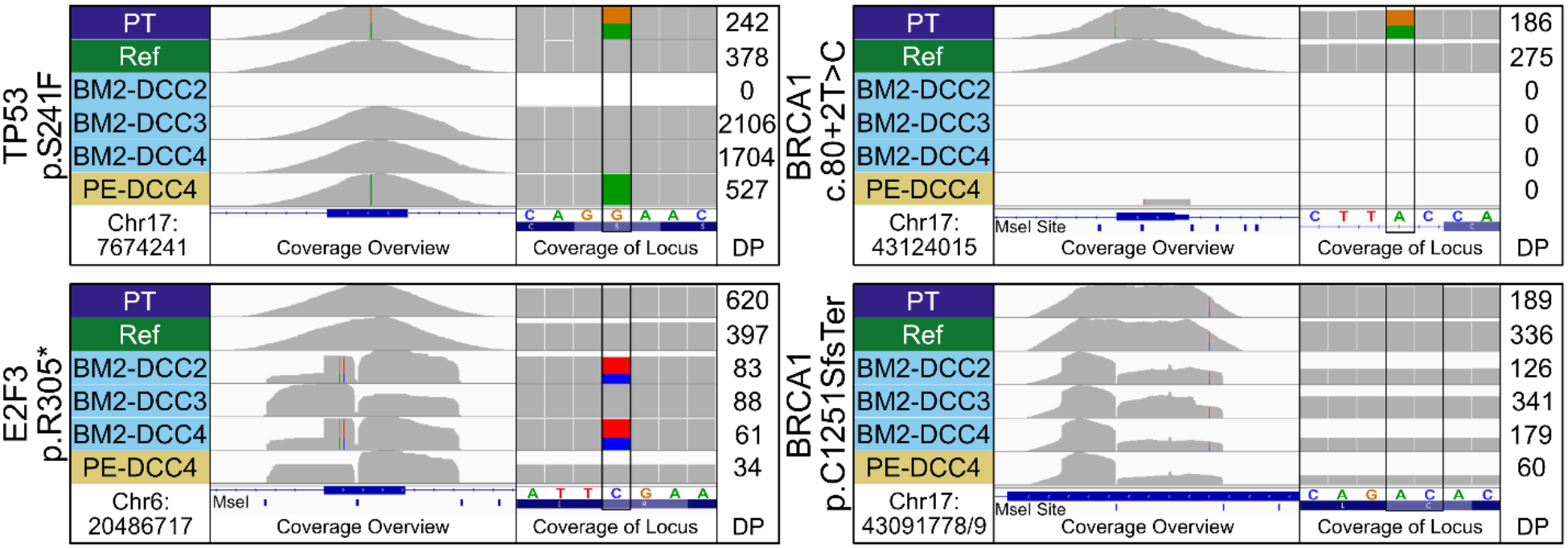
Coverage profiles of selected genomic loci in representative samples. Representative exon-level and locus-specific sequencing depth distribution profiles. DP – values denote total read depth at the specific genotype call. The positions of the MseI restriction sites are indicated below the depicted exons.

**Figure S4.**
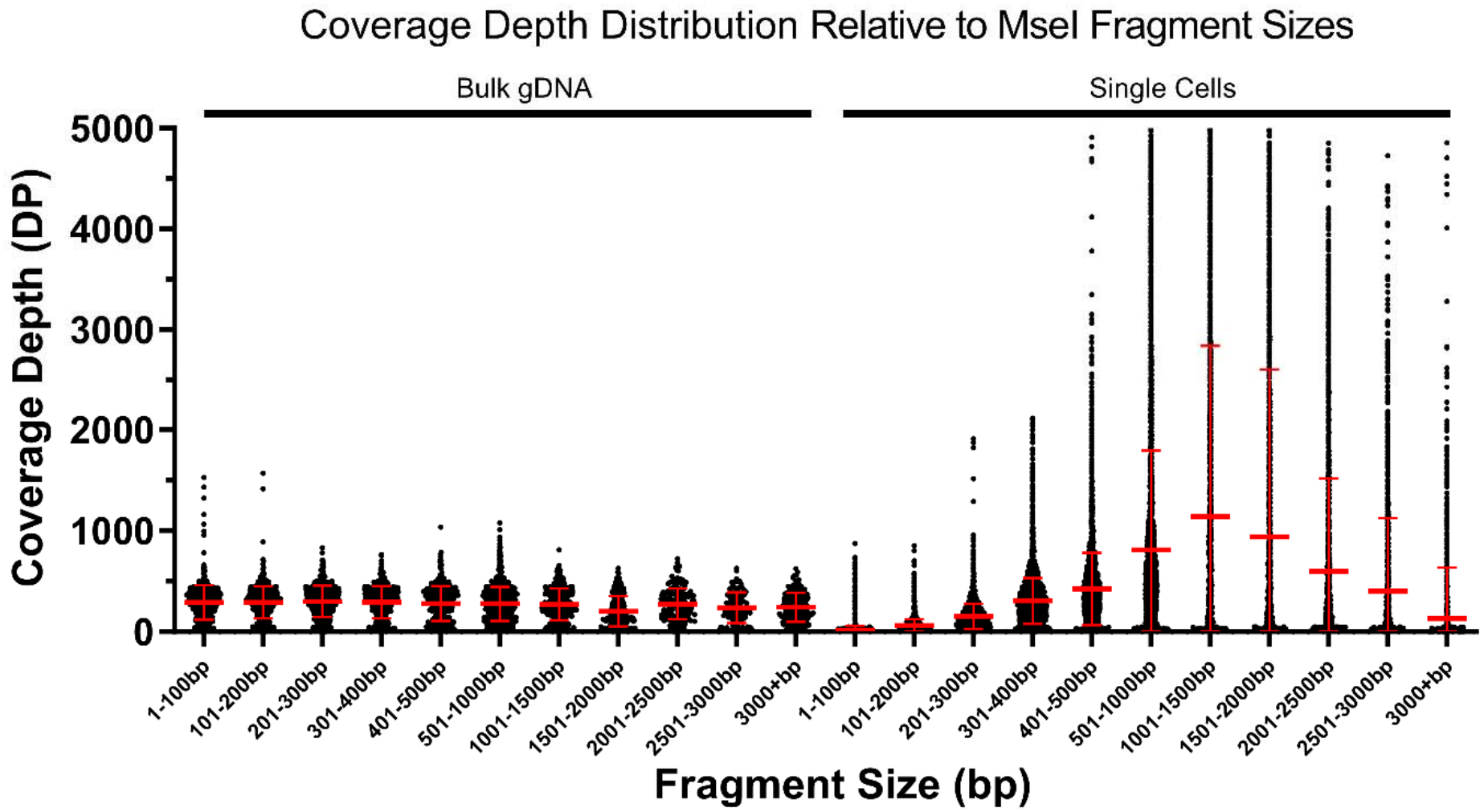
Coverage bias is introduced by the WGA process. SNV coverage depth across MseI restriction fragment stratified by size in bulk DNA (left) and single-cell WGA samples (right). Bulk DNA showed uniform coverage across all fragment size ranges, whereas WGA products displayed size-dependent coverage bias, with reduced representation of SNVs located on short (<300bp) and long (>3000bp) fragments. Red bars indicate the mean (± standard deviation) SNV coverage depth for variants mapping to MseI restriction fragments within the selected size ranges.

