## Supplementary Methods for "Monitoring cancer evolution under iatrogenic selection pressure by cellular liquid biopsy"

**Patient characteristics, clinical staging, and sample processing.**

The patient, treated at the Comprehensive Cancer Center Regensburg, provided written informed consent for comprehensive molecular profiling of primary tumor tissue, circulating tumor cells (CTCs), disseminated cancer cells (DCCs), pleural effusion (PE) specimens and matched germline controls, as well as for review of clinical data including pathology and treatment history (ethics approval ID: 18-948-101). All procedures were conducted in accordance with the Declaration of Helsinki.

Treatment history and clinical characteristics are summarized in Table S1 and in Figure 1. Briefly, the patient was diagnosed with 4.1cm, non-metastatic, invasive ductal carcinoma of no special type in the right breast. Neoadjuvant chemotherapy consisting of epirubicin plus cyclophosphamide followed by paclitaxel was administered between months 2 (Timepoint 3, TP3) and 6 (TP7) after diagnosis. Surgical resection of the primary tumor with axillary lymph node dissection was performed 8 months after diagnosis (TP9). Histopathological assessment classified the tumor as luminal B–like (ER IRS 4, PR IRS 3, HER2 1+, Ki67 75%). Adjuvant therapy with abemaciclib, letrozole and goserelin was administered between months 10 and 15 after diagnosis (TP11 and TP15, respectively). During adjuvant treatment, a 1.4 cm liver metastasis was detected. Biopsy of the metastatic lesion 16 months after primary diagnosis (TP17) revealed a triple-negative phenotype (ER-negative, PR-negative, HER2-negative); however, the lesion consisted of approximately 50 cells making the immunohistochemical assessment insufficient to confidently confirm a phenotype transformation into triple-negative breast cancer. Between months 17 and 21 after diagnosis, the patient received systemic treatment with (i) nab-paclitaxel plus atezolizumab, (ii) carboplatin and (iii) olaparib. The patient died 23 months after initial diagnosis (TP24). Bone marrow (BM) aspirates were collected at months 1 and 8 after diagnosis (before and after neoadjuvant therapy), yielding 0 and 5 DCCs, respectively. Peripheral blood samples were collected at months 17, 19, 21 and 22 during metastatic treatment, yielding 476, 804, 246, and 8,783 CTCs per 7.5 mL of blood, respectively. In addition, one pleural effusion sample was obtained at month 18 (TP19), prior to initiation of palliative treatment.

In total, 38 breast cancer–related specimens were analyzed, including 2 bulk DNA samples and 36 single-cell WGA products. The cohort included one bulk PBMC-derived germline DNA sample and 1 single PBMC isolated from peripheral blood serving as germline controls. Three single-cell WGA samples yielded low-quality results and were eliminated from the analysis, bringing the total number of samples used in the study to 35. A complete overview of all specimens analyzed is provided in Table S1. Bulk genomic DNA isolate extracted from the formalin-fixed paraffin-embedded (FFPE) primary tumor tissue was obtained from the Institute of Pathology, University Hospital Regensburg.

**Single-cell detection protocols (DCCs, CTCs).**

DCC enrichment and detection

BM aspirates were obtained from the anterior iliac crest. One PE sample was additionally collected. BM and PE specimens were processed on the day of sampling. Mononuclear cells were enriched by density gradient centrifugation. Samples were washed in Hanks’ Balanced Salt Solution (HBSS; Anprotec, Cat. No. AC-BS-0022) and subjected to Percoll density gradient centrifugation (density 1.077 g/mL; (Thermo Fisher Scientific, Cat. No. 10607095)) according to established protocols ^1^. The mononuclear cell fraction was collected and transferred onto adhesion slides (0.25 × 10⁶ cells per 227 mm²; (Epredia, Cat. No. X2XER203B), air-dried overnight and stored at −20 °C until staining. When available, 2 × 10⁶ cells (two slides) were screened per timepoint. Immunocytochemical detection of DCCs was performed using either the monoclonal antibody A45-B/B3 recognizing cytokeratins 8, 18 and 19 (Micromet / AS Diagnostics, Cat. No. FAR002A). An isotype-matched IgG1 κ antibody (clone MOPC-21; Sigma Aldrich, Cat. No. M9269) was used as negative control.

Signal detection was performed using the ZytoChem Plus (AP) Polymer Kit (Zytomed Systems, Cat. No. ZUC077) with 5-bromo-4-chloro-3-indolyl phosphate/nitroblue tetrazolium (BCIP/NBT; Bio-Rad, Cat. No. 1706432) as a substrate. Slides were evaluated by brightfield microscopy. DCC enumeration was performed blinded to the clinical timepoint. Inter-observer validation was performed by independent investigators.

CTC enrichment and detection (CellSearch®)

Peripheral blood samples were collected between December 2023 and March 2024. Blood was processed within 96 hours of collection. For each timepoint, 7.5 mL of peripheral blood was collected into CellSave® Preservative Tubes (Menarini Silicon Biosystems, Cat. No. CS0013). CTC enrichment and enumeration were performed using the CellSearch Circulating Tumor Cell Kit (Menarini Silicon Biosystems, Cat. No. CS0009) according to the manufacturer’s instructions and as previously described ^2^).

CTCs were enriched by immunomagnetic separation using ferrofluid nanoparticles coated with anti-EpCAM antibody (clone VU1D9). Enriched cells were fluorescently labeled with antibodies directed against cytokeratins 8/18 (clone C11) and cytokeratin 19 (clone A53-B/A2), as well as against CD45 (clone HI30). Nuclear staining was performed using DAPI (4′,6-diamidino-2-phenylindole). Cells exhibiting a CK⁺/CD45⁻/DAPI⁺ staining pattern were classified as CTCs, whereas CK^±^/CD45⁺/DAPI⁺ cells were classified as leukocytes. CTC enumeration was performed blinded to the clinical timepoint.

**DNA extraction, whole-genome amplification, and quality control.**

Single-cell isolation

Single cells were identified according to immunophenotypic criteria and isolated by manual micromanipulation. CTCs were defined as CK⁺/CD45⁻/DAPI⁺ events, whereas leukocytes were defined as CK^±^/CD45⁺/DAPI⁺ events. DCCs, defined as CK^+^ cells in BM and PE samples, and corresponding CK^−^ control cells were collected from adhesion slides following the immunohistochemistry staining described above.

Manual micromanipulation

Cells recovered from the CellSearch cartridge were washed in 1× phosphate-buffered saline (1x PBS) and transferred onto bovine serum albumin (BSA)-coated slides, whereas cells deposited on adhesion slides were overlaid with 1x PBS and isolated directly from the slide surface. Individual cells were classified as CTCs, DCC or control cells based on their staining pattern (see above) using an inverted microscope (Olympus, Model: BX43F, Cat. No. BX43TF) and isolated using a micromanipulator system (Eppendorf, Model: Patchman NP2, Cat. No. 1049027296).

Single cells were collected in 1 µL of 1× PBS and immediately subjected to whole-genome amplification (WGA) as previously described (see below).

DNA extraction from bulk material

Genomic DNA from an FFPE specimen of primary tumor tissue was extracted using the QIAamp DNA FFPE Tissue Kit (Qiagen, Cat. No. 56404) according to the manufacturer’s protocol. Genomic DNA from peripheral blood mononuclear cells (PBMCs) was isolated using the DNeasy Blood & Tissue Kit (Qiagen, Cat. No. 69504). For each extraction, 3 × 10⁶ PBMCs were used as input material. The concentration and purity of the extracted DNA were assessed using a NanoDrop UV-Vis Spectrophotometer (Thermo Fisher Scientific, Model: NanoDrop 2000c UV-Vis, Cat. No. ND-2000C) and a Qubit fluorometer (Invitrogen, Model: Qubit 3.0, Cat. No. Q33216). The concentration of the double-stranded DNA (dsDNA) fraction was additionally quantified using either the Qubit dsDNA BR or the 1X dsDNA BR kits (Invitrogen-Thermo Fisher Scientific, Cat. No. Q32853 and Q33266).

Whole-genome amplification (WGA) and quality control

Single-cell WGA was performed using a deterministic adapter-linker PCR–based method as previously described ^3,4^. The WGA workflow consists of proteolytic digestion, restriction digestion, adapter pre-annealing, and ligation, followed by PCR amplification. Importantly, in this WGA approach, the single-cell genome is deterministically fragmented with the restriction enzyme MseI before adapter ligation and amplification. This generates a consistent and reproducible representation of single-cell DNA across samples. However, as a consequence, long (>3,000 bp) and short (<100 bp) MseI restriction fragments are underrepresented in the resulting WGA products, owing to the limited processivity of the DNA polymerase used during PCR and loss of short fragments during post-WGA processing, respectively (Fig. S4). Given the limited DNA content of single CTCs and DCCs and the irreversible nature of WGA, no technical replicates were performed. No-template controls (NTCs) were included in each WGA run to monitor contamination. WGA product quality was assessed using a validated 4-plex endpoint PCR assay (WGA-QC) enabling calculation of the genome integrity index (GII) as previously described ^5^. Only samples with GII ≥ 3 were included in downstream analyses.

Pre-processing of WGA products for NGS

To remove WGA adapter sequences prior to next-generation sequencing (NGS) library preparation, WGA products underwent enzymatic pre-processing. Briefly, 5 µL of WGA product was subjected to dsDNA synthesis. To this purpose, DNA templates were supplemented with 0.1 µL dNTP mix (10 mM each), 0.1 µL universal primer (100 µM; 5′-AGTGGGATTCCGCATGCTAGT-3′), 0.05 µL DNA polymerase (5 U/µL; Roche Diagnostics, Cat. No. 11759060001), 0.1 µL Expand Buffer 2 (Roche Diagnostics, Cat. No. 11759060001) and 0.65 µL PCR-grade water. Samples were incubated at 68 °C for 2 hours to convert single-stranded WGA amplicons into dsDNA. Adapter sequences were subsequently cleaved by digestion with 0.12 µL Tru1I restriction enzyme (50 U/µL; Thermo Fisher Scientific, Cat. No. ER0983) at 65 °C for 3 hours. Digestion products were purified using magnetic bead–based cleanup with either AMPure XP beads (Beckman Coulter, Cat. No. A63881) or KAPA HyperPure Beads (Roche Diagnostics, Cat. No. 8963851001) at a bead-to-sample ratio of 1.8× according to the manufacturer’s instructions.

**Targeted sequencing of DNA**

Sequencing of bulk DNA and single-cell WGA products with the scIMPACT assay.

Targeted sequencing was performed using a custom hybridization-based assay termed single-cell Integrated Mutation Profiling of Actionable Cancer Targets (scIMPACT), which was adapted and optimized to facilitate the SNV and CNV profiling of single-cell WGA products. The panel design (IMP2.0) was based on the FDA-authorized MSK-IMPACT assay (MSK-IMPACT FDA Decision Summary DEN170058 ^6^), albeit it was expanded to include the coding sequence of additional 12 genes and selected pharmacogenomics (PGX) markers implicated with drug metabolism in breast, lung, prostate cancer and malignant melanoma. The PGX makers were selected from the PharmGKB knowledge base. The final design encompasses the complete coding sequences of 480 cancer-associated genes and single-nucleotide polymorphism loci across 120 genes, comprising a total of 8,240 capture targets spanning ~2 Mb of genomic sequence. The list of targeted genes is summarized in Table S5.

Sequencing libraries were prepared using the KAPA HyperPrep Kit (Roche Diagnostics, Cat. No. 07 962 363 001) and KAPA HyperCap v3.3 reagents (Roche Diagnostics, KAPA Universal Adapter, Cat. No. 09 063 781 001) according to the manufacturer’s instructions. For all samples, including bulk genomic DNA and pre-processed single-cell WGA products, 110 ng of double-stranded DNA was used as standardized input to ensure uniform library complexity. DNA was acoustically sheared to a target fragment peak size of 200–250 bp, followed by end repair, A-tailing, adapter ligation, and 7–8 cycles of pre-capture PCR incorporating KAPA unique dual indices (Roche Diagnostics, KAPA UDI Primer Mixes 1-96 and 97-192, Cat. No. 09 134 336 001and 09 329 838 001). Amplified libraries were purified twice using magnetic KAPA HyperPure beads (Roche Diagnostics, Cat. No. 8963851001) and pooled in equimolar amounts to generate pre-capture library pools comprising 20-21 libraries each, with a total 2 µg DNA per pool. Hybridization-based target enrichment was performed to generate post-capture library pools using custom-designed biotinylated RNA probes (Roche Diagnostics, Cat. No. 09053069001) with 1.5 µg of pooled library DNA as input, followed by 14 cycles of post-capture PCR and two additional bead purification steps. To facilitate genome-wide CNV analysis through retention of off-target reads, post-capture libraries were supplemented with the corresponding pre-capture pool at an 85:15 ratio prior to sequencing. Paired-end sequencing (2 × 150 bp) was conducted on an Illumina NovaSeq 6000 instrument, targeting a minimum of 15 million paired-end read per sample. Bulk DNA samples achieved a mean coverage of 578× (standard deviation; SD 278×), whereas whole-genome amplified single-cell libraries achieved a mean unique coverage of 551× (SD 241×).

Targeted sequencing of bulk tumor and germline DNA (Illumina TruSight™ Oncology 500).

DNA sample derived from FFPE primary tumor tissue was processed using the TruSight™ Oncology 500 (TSO500) assay (Illumina, Cat. No. 20130528) according to the manufacturer’s instructions. Library preparation, hybrid-capture target enrichment, and indexing were performed using the recommended protocol for FFPE-derived DNA input amounts. Sequencing was performed on an Illumina platform (Illumina 6000Dx, Cat. No. 20068232).

Sequencing quality metrics met manufacturer-recommended thresholds. On-target alignment rate and uniformity of coverage were within acceptable ranges as defined by the TSO500 performance guidelines. Coverage of exonic nucleotides relevant for mutation and tumor mutational burden (TMB) analysis reached 99.5% at ≥50 UMI reads, surpassing the recommended 90% threshold.

Primary data processing, alignment to the human reference genome (GRCh38/hg38), small variant calling, copy-number alteration detection and tumor mutational burden (TMB) estimation, as well assessment of the microsatellite instability (MSI) were performed using the DRAGEN Bio-IT Platform with the dedicated TSO500 analysis module, using default parameters unless otherwise specified. Variants were reported when supported by at least 50 UMI reads and a mutant allele fraction (MAF) of ≥5%. TMB was calculated as the number of somatic, coding base substitution and indel mutations per megabase (Mb) of targeted genome territory as defined by the assay. Microsatellite instability (MSI) was assessed using the Illumina TSO500 assay by evaluating length variability across a panel of 130 A/T homopolymer microsatellite loci and comparing their distribution to a panel of normal samples using the DRAGEN TSO500 bioinformatics pipeline. MSI status was classified as high when ≥10% of evaluated microsatellite loci (with a minimum of 40 loci assessed) exhibited a length distribution significantly different (Jensen–Shannon distance ≥ 0.1, p-value ≤ 0.01) from that observed in the reference normal dataset.

Targeted sequencing of BRCA1/2 and HRD assessment (QIAGEN QIAseq Targeted DNA IO Panel).

Tumor DNA was additionally analyzed using the QIAGEN QIAseq Targeted DNA IO Panel to sequence the coding regions of *BRCA1* and *BRCA2* and to assess homologous recombination deficiency (HRD). Library preparation and target enrichment were performed according to the manufacturer’s protocol for FFPE-derived DNA input, followed by sequencing on an Illumina platform. Variants were reported at a MAF detection threshold of ≥5%.

HRD was inferred from genome-wide copy-number and allelic imbalance profiles derived from the panel’s SNP backbone. The HRD score was calculated as the sum of loss of heterozygosity, telomeric allelic imbalance and large-scale state transitions, and samples with an HRD score ≥56 were classified as HRD-positive.

Sequencing quality metrics met manufacturer-recommended thresholds. Median sequencing depth reached 528 UMI-supported reads, and 98.52% of the targeted regions were covered, exceeding the recommended minimum coverage threshold of 95%.

**Bioinformatic Analysis for SNV, Indel, and CNV Detection in the scIMPACT workflow.**

Paired-end FASTQ files from patient-derived samples and DNA derived from the GM12878 cell line, used as a normal reference, were used as input for variant calling.

SNV and insertion/deletion (indel) analyses were performed using the human reference genome GRCh38/hg38 from GATK resource bundle, with variant calling restricted to the target regions defined in the corresponding IMP2.0 capture BED file.

SNV and Indel Calling

Quality of raw sequencing data was assessed using FastQC v0.11.9, evaluating per-base sequence quality, GC content, adapter contamination, duplication rates, length distribution, and overrepresented sequences. Adapter trimming and quality filtering were performed using BBDuk (BBTools suite v2.0.13), wherein bases with Phred quality scores <10 were trimmed from read ends, and the first three base pairs of each read were hard clipped to mitigate positional bias. Reads shorter than 50 bp after trimming were discarded.

Subsequently, to ensure high data quality, reads mapping to microbial, fungal, or common laboratory contaminant sequences were removed and only reads that were mapping exclusively to the human reference genome were retained for downstream analyses. Where appropriate, overlapping paired-end reads were merged to reduce sequencing error rates.

High-quality trimmed decontaminated reads were aligned to GRCh38/hg38 using BWA-MEM (v0.7.17). PCR duplicates in the aligned reads were marked with Picard MarkDuplicates (v2.21.8). Alignment quality metrics were evaluated using Qualimap (v2.2.2d) and samtools (stats, flagstat).

Subsequently, base quality score recalibration (BQSR) was performed using GATK (v4.2.6.1). Known polymorphic sites from dbSNP v138, HapMap v3.3, 1000 Genomes Phase 1 high-confidence SNPs, and Axiom Exome Plus were used to model and detect systematic errors. Recalibration values were estimated with GATK BaseRecalibrator and applied, corrected using GATK ApplyBQSR producing final analysis-ready BAM files. Diagnostic plots were generated using GATK AnalyzeCovariates to evaluate recalibration performance.

Somatic as well as germline SNVs and small indels were identified using multiple variant callers, including Mutect2 (GATK v4.2.6.1), FreeBayes (v1.3.6), and Strelka (v2.9.10), to maximize detection sensitivity and robustness. Mutect2 was run in tumor–normal paired mode using gnomAD as a germline resource and a Panel of Normals (PoN) constructed according to GATK best practices.

Variant call format (VCF) files from all callers were merged using bcftools merge to generate a unified VCF file. Variants were left-aligned and trimmed to standardize representation, multi-nucleotide polymorphisms were decomposed using vt decompose, and redundant loci were normalized and consolidated.

Functional annotation was performed using ANNOVAR (2020-06-08), Ensembl Variant Effect Predictor (VEP v105), and SnpEff (v5.0). Annotation databases included ClinVar, COSMIC, dbSNP, gnomAD, and population allele frequency datasets. To incorporate read-level quantitative support for the SNV and indels, bam-readcounts was used, which were incorporated into the final VCF files using VAtools.

Variant prioritization focused on high-confidence somatic events was facilitated by the Bayesian classification workflow (described below). Germline variants were excluded using matched normal or blood-derived DNA. Retained variants included those with *Necessary* (high confidence) or *Warranted* (medium confidence) classification as defined by the COMPASS workflow (see below), particularly protein-altering variants predicted as “probably damaging” or “possibly damaging,” and variants with known or reported oncogenic associations.

Bayesian classification workflow for variant prioritization.

To identify and prioritize variants supported by high variant allele frequency (VAF) and established or predicted pathogenic relevance, we implemented a Bayesian classifier–based workflow adapted from a previously described framework ^7^. This Bayesian training approach allows to compute the balancing factors for regularization instead of optimizing them using cross validation, which is especially suitable for smaller data sets with class imbalances.

The classifier was generated using scIMPACT sequencing data derived from a reference cohort of 28 samples generated from well-characterized breast cancer cell lines (BT-474, MDA-MB-453, BT-549 and ZR-75.1). Variants annotated as present in the respective cell lines in at least one of three curated databases - Sanger Cell Model Passport, DepMap and COSMIC Cell Lines Project - were considered true-positive events for training. The corresponding data were downloaded from DepMap on 26 October 2020, from the COSMIC Cell Lines Project on 6 July 2021, and from the Sanger Cell Model Passport on 11 September 2022.

Each of the cell lines represents 25% of the sequenced sample cohort, wherein two of the cell lines (BT-474 and MDA-MB-453) contain a higher number of variants with positive labels (expected/true-positive events). The internal validation strategy consisted of using data from two cell lines – one with a high and one with a low number of expected mutations – for training, and the remaining two cell lines for validation.

Approximately 4% of the sequencing output data were labeled as true-positive events, resulting in a strong class imbalance that makes training the classifier challenging. Reweighted class contributions were applied during training to emphasize the true-positive class and counter the class imbalance. Consequently, each variant was included with a probability of 90% if it had a positive label (expected mutation) and 20% if it had a negative label (unexpected mutation). This resampling strategy increased the proportion of positive events in the training set to about 17%. No resampling was applied to the validation data. All feature processing steps were performed in Python (version [3.10.13]), classifier training and classifier application are Java libraries (version [23.0.2-internal 2025-01-21]).

The classifier integrates fifteen features capturing variant abundance (reflected by variant allele frequency; VAF), sample type (bulk DNA, single cell or pooled cells), and the following annotation scores: M-CAP_rankscore, FATHMM_converted_rankscore, fathmm-XF_coding_rankscore, MutationAssessor_score, MPC_score, phyloP30way_mammalian_rankscore, GenoCanyon_score, BayesDel_addAF_score, PolyPhen-2_HDIV_score, SIFT_score, SIFT4G_converted_rankscore and LRT_score. No additional feature preprocessing steps, such as scaling or normalization, were necessary, as all features have a suitable range already. Variant annotation was performed using the dedicated tools described above.

The Bayesian model generated a continuous probability score hereby referred to as the ***Clinical Oncology Mutation Pathogenicity and Actionability Scoring System*** (COMPASS) ranging from 0 to 1, corresponding to increasing confidence that a variant represents a true somatic event with pathogenic relevance. COMPASS classification tiering consists of four categories referring to decreasing levels of confidence: *Necessary, Warranted, Speculative, and Exempt*.

Based on the training dataset, COMPASS thresholds were defined as ≥0.752 for *Necessary* or high-confidence variants (specificity 99.42%, sensitivity 67.44%, precision 90%) and ≥0.617 for *Warranted* or medium-confidence variants (specificity 98.93%, sensitivity 78.39%, precision 85%). Variants scoring below 0.617 were classified as *Speculative* or low-confidence variants, whereas variants without a COMPASS score were classified as *Exempt* because they did not meet the minimum requirements for classification. Exempt variants included loci with insufficient total reads depth (DP < 20) or insufficient alternate-allele support (AD < 10 and/or VAF < 0.1), consistent with inadequate coverage or absence of a confidently detectable mutation, respectively. Sensitivity, specificity and precision were calculated using standard definitions (TP/(TP+FN)), (TN/(TN+FP)) and TP/(TP+FP), respectively.

The classification workflow was implemented using proprietary in-house code and is available from the corresponding author upon reasonable request and subject to institutional data-sharing agreements.

Copy Number Variant (CNV) Analysis

Copy number variant (CNV) analysis was performed using off-target sequencing reads, i.e. reads mapping to the genomic loci located outside of the target region defined by the respective panel BED file (IMP2.0 design). For the purpose of CNV profiling, the human genome was divided into non-overlapping bins of 500 kb. CNV calling was conducted independently for each sample. Following adapter trimming and removal of low-quality bases using BBDuk (v38.84), decontamination was performed using BioBloomTools (v2.0.13) to eliminate non-human reads originating from microbial or fungal DNA, as well as potential contaminants from water or reagents. Remaining reads were aligned to the human reference genome (GRCh38/hg38) using BWA-MEM (v0.7.17). Aligned reads were counted per genomic bin, filtered, corrected for GC-content and mappability, normalized, and segmented using the Bioconductor package QDNAseq (v1.34.0). The same package was used to generate log2 ratio CNV profiles.

Integrated plots combining CNV data from multiple samples were generated to provide a comprehensive visualization of genomic alterations outputted by the QDNAseq package. The percentage of aberrant genome (PAG) was calculated as: PAG=100×s_a_ / s_g_, where s_g_ represents the total size of the binned genome and s_a_ the cumulative size of aberrant regions, both measured in base pairs. Quality control thresholds were applied to ensure reliability of CNV profiles. Samples with fewer than 100,000 mapped off-target reads were excluded because QDNAseq cannot generate reliable CNV profiles from such limited amounts of data. Additionally, the measured standard deviation (σ̂Δ), calculated using an estimator based on von Neumann’s mean square successive differences, was used as a noise metric. Profiles with σ̂Δ > 0.5 were excluded. Profiles with σ̂Δ > 0.2 were retained but interpreted with caution, as noise at this level may partially obscure true signal.

The combined SNV/indel and CNV analysis framework enabled comprehensive detection and interpretation of genomic alterations in tumor samples within a clinically oriented sequencing workflow.

Data availability.

Raw sequencing data and processed variant call files are available from the corresponding author upon reasonable request, subject to institutional and ethical regulations governing patient-derived genomic data. Access will require approval by the responsible ethics committee and execution of a data transfer agreement in accordance with applicable data protection regulations.
