## Supplementary Figure 1 for "Monitoring cancer evolution under iatrogenic selection pressure by cellular liquid biopsy"

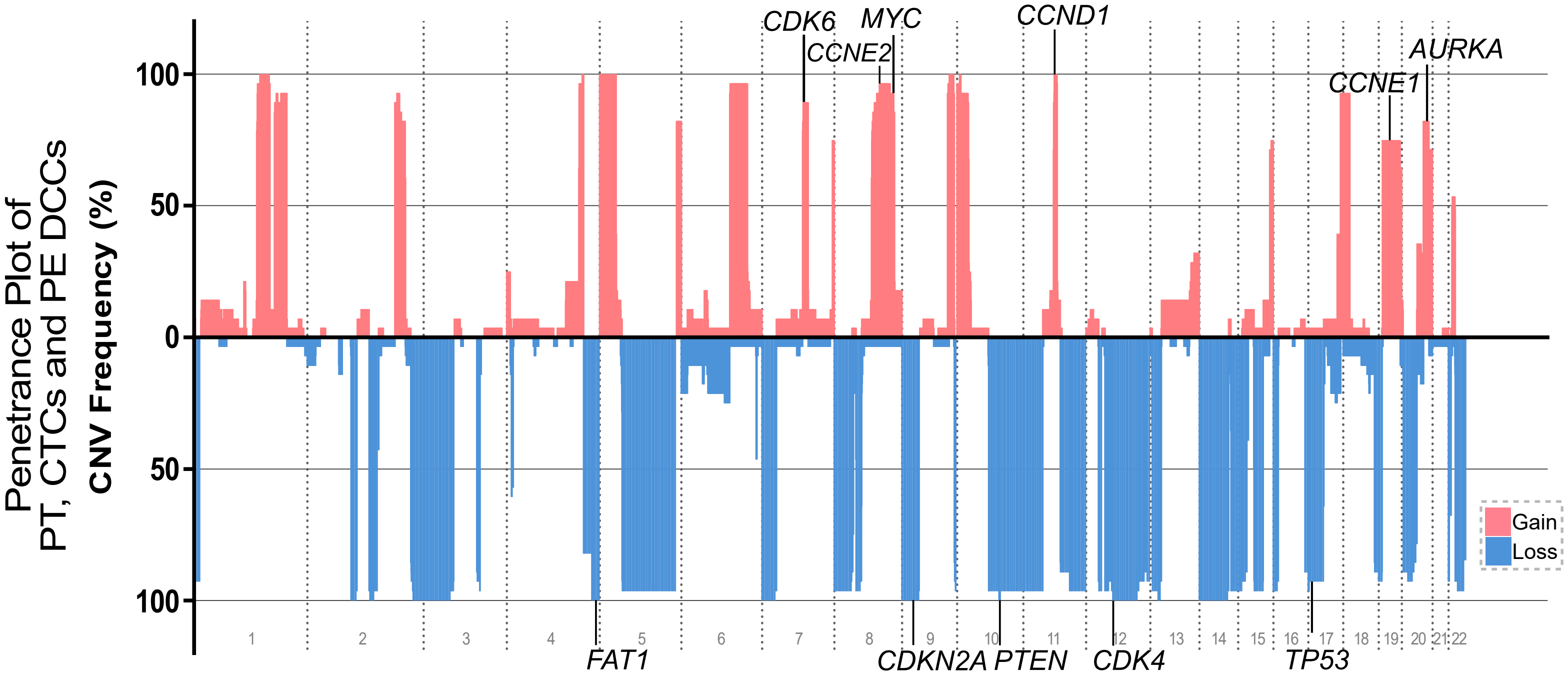

Penetrance Plot of
PT, CTCs and PE DCCs
CNV Frequency (%)
100
50
0
50
100
CDK6
CCNE2
MYC
CCND1
CCNE1
AURKA
FAT1
CDKN2A
PTEN
CDK4
TP53
Gain
Loss
