## Supplementary Figure 2 for "Monitoring cancer evolution under iatrogenic selection pressure by cellular liquid biopsy"

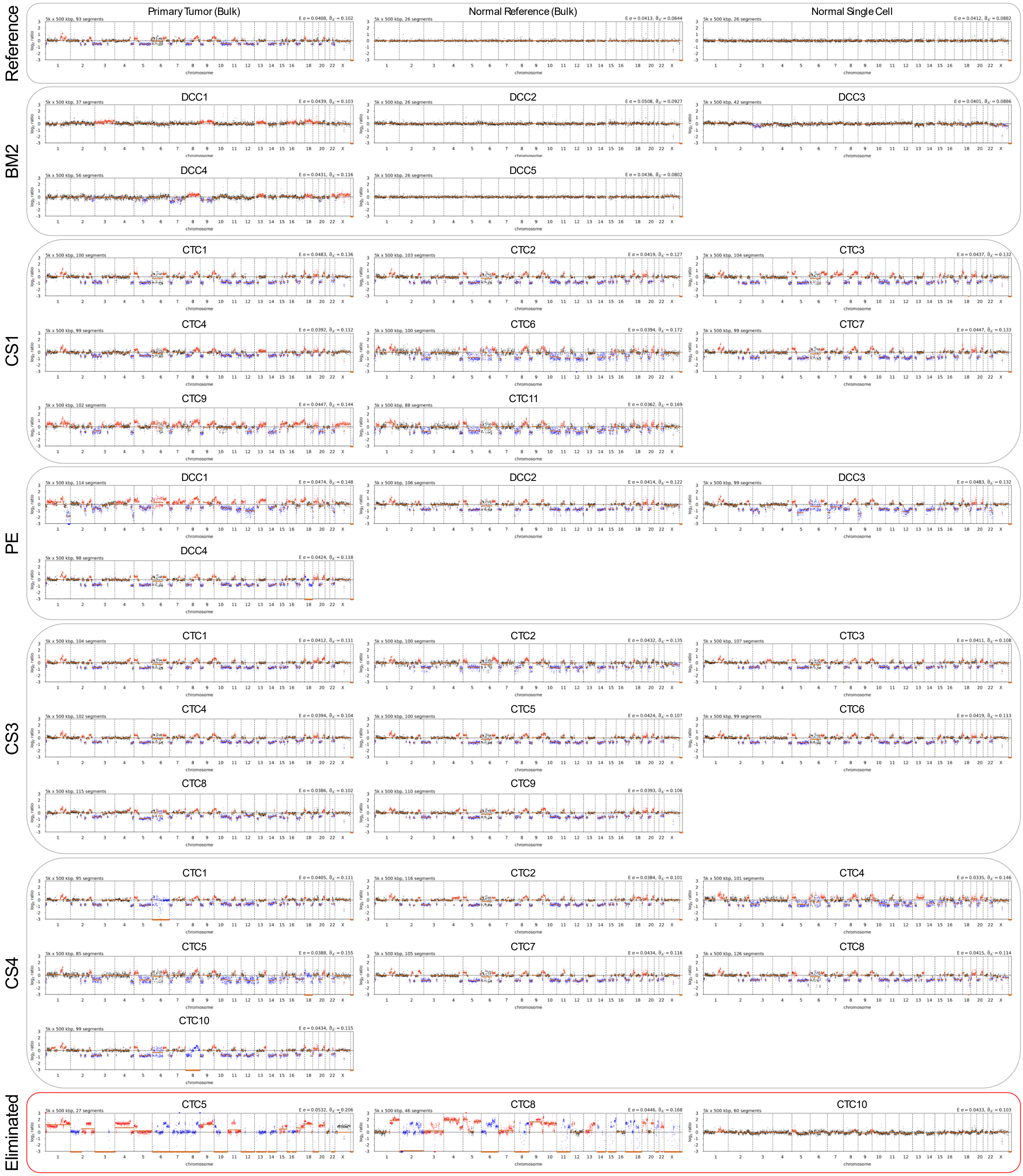

Reference
Primary Tumor (Bulk)
Normal Reference (Bulk)
Normal Single Cell
BM2
DCC1
DCC2
DCC3
DCC4
DCC5
CS1
CTC1
CTC2
CTC3
CTC4
CTC6
CTC7
CTC9
CTC11
PE
DCC1
DCC2
DCC3
DCC4
CS3
CTC1
CTC2
CTC3
CTC4
CTC5
CTC6
CTC8
CTC9
CS4
CTC1
CTC2
CTC4
CTC5
CTC7
CTC8
CTC10
Eliminated
CTC5
CTC8
CTC10
chromosome
