## Supplementary Figure 3 for "Monitoring cancer evolution under iatrogenic selection pressure by cellular liquid biopsy"

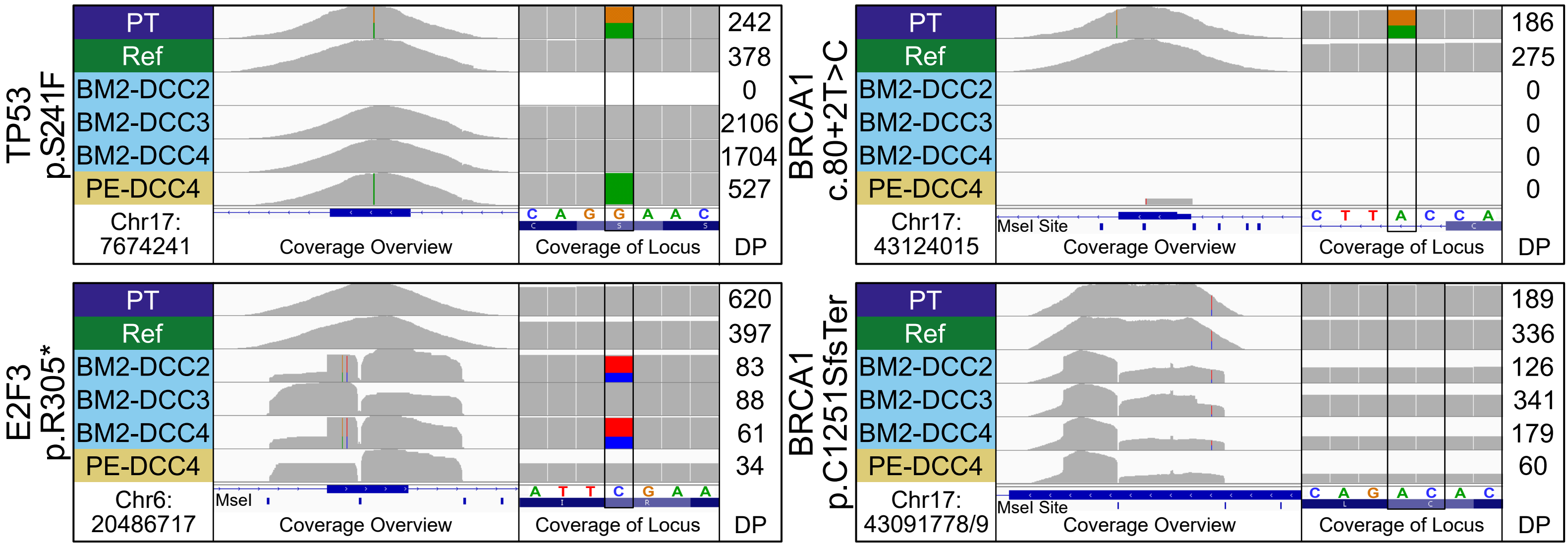
TP53
p.S241F
PT
Ref
BM2-DCC2
BM2-DCC3
BM2-DCC4
PE-DCC4
Chr17:
7674241
Coverage Overview
Coverage of Locus
DP
242
378
0
2106
1704
527
C A G G A A C
BRCA1
c.80+2T>C
Chr17:
43124015
MseI Site
186
275
0
0
0
0
C T T A C C A
E2F3
p.R305*
Chr6:
20486717
MseI
620
397
83
88
61
34
A T T C G A A
BRCA1
p.C1251SfsTer
Chr17:
43091778/9
189
336
126
341
179
60
C A G A C A C
