## Supplementary Figure 4 for "Monitoring cancer evolution under iatrogenic selection pressure by cellular liquid biopsy"

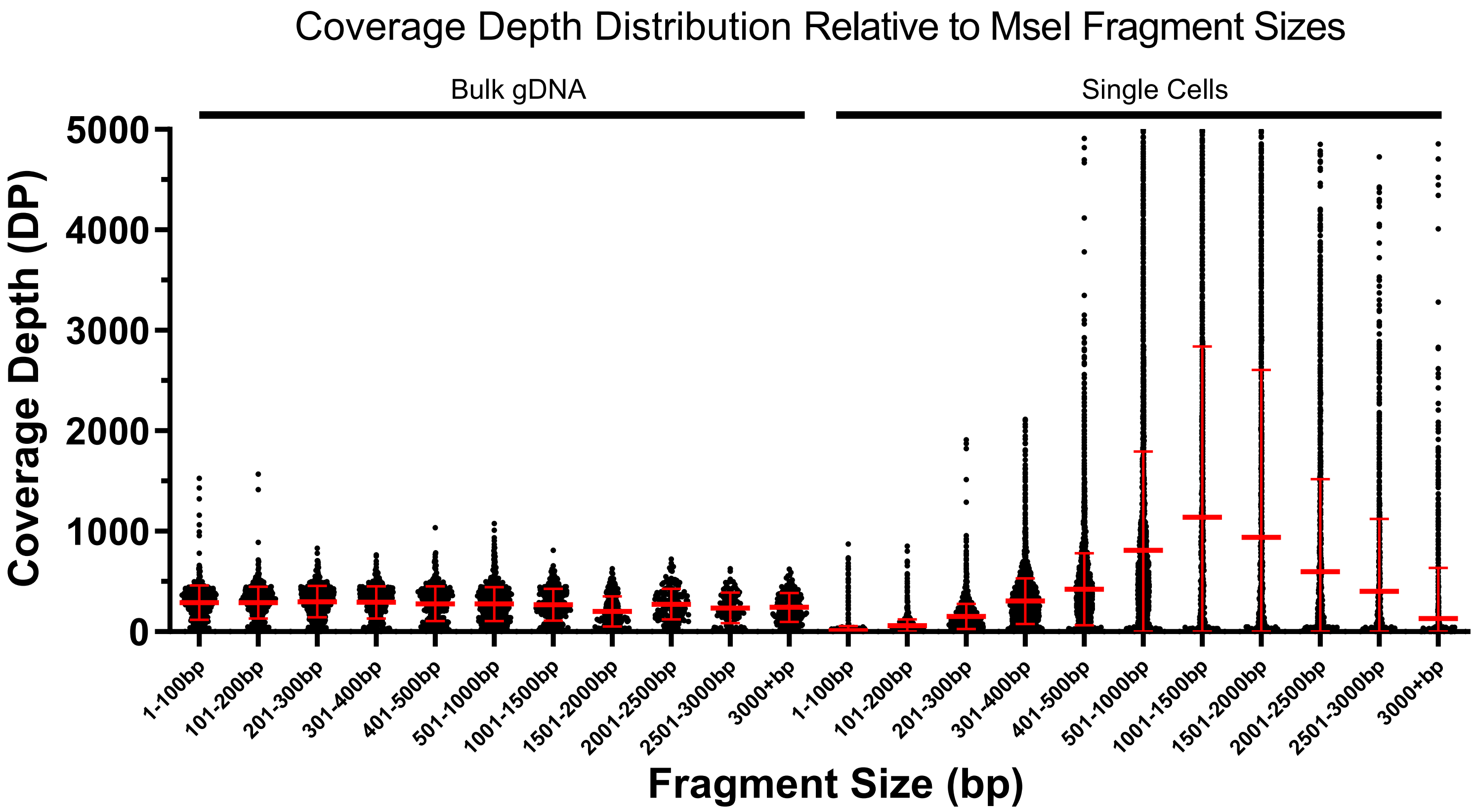
Coverage Depth Distribution Relative to MseI Fragment Sizes
Bulk gDNA
Single Cells
Coverage Depth (DP)
5000
4000
3000
2000
1000
0
1-100bp
101-200bp
201-300bp
301-400bp
401-500bp
501-1000bp
1001-1500bp
1501-2000bp
2001-2500bp
2501-3000bp
3000+bp
1-100bp
101-200bp
201-300bp
301-400bp
401-500bp
501-1000bp
1001-1500bp
1501-2000bp
2001-2500bp
2501-3000bp
3000+bp
Fragment Size (bp)
